# Genomic Evolution of SARS-CoV-2 Delta Variants Pre- and Post-Omicron Emergence using Alignment-free Machine Learning models

**DOI:** 10.64898/2026.02.20.706927

**Authors:** Sathish Sankar, Kaushika Anandharaman, Pradeesh Selvam, Aswini Jayaraman, Deepak Jayakumar, Pachamuthu Balakrishnan, Marie Larsson, Vijayakumar Velu, Raju Sivadoss, Esaki M. Shankar

## Abstract

The SARS-CoV-2 Delta variant (B.1.617.2), initially classified as a variant of concern due to its enhanced transmissibility and vaccine-escape mutations, underwent further genomic changes following the emergence of the Omicron variant (B.1.1.529). This study investigates the genomic differences in Delta variant spike gene sequences collected before and after the emergence of Omicron. A total of 190 sequences were analyzed using an alignment-free approach incorporating k-mer-based feature extraction and machine learning models, including convolutional neural networks (CNN), K-means clustering, and random forest classification. The random forest model achieved 93% accuracy, with significant F1 scores, effectively distinguishing the two Delta variant groups. Comparative analysis revealed 157 persistent mutations and four vanished mutations in the post-Omicron group. Cluster analysis showed notable shifts, indicating stable yet evolving genomic patterns over time. The study demonstrates the advantage of alignment-free methods in detecting subtle sequence variations that alignment-based approaches may overlook. These findings enhance our understanding of SARS-CoV-2 evolution and provide a framework for identifying key genomic signatures relevant to public health. The methodology and insights gained offer potential applications in variant surveillance, vaccine design, and viral evolutionary studies, supporting preparedness for future SARS-CoV-2 variant emergence.

## 1 Introduction

The WHO estimates suggest that the severe acute respiratory syndrome coronavirus 2 (SARS-CoV-2) pandemic recorded over 777 million cases with >7 million deaths globally[1]. Active extensive surveillance of COVID-19 has been discontinued due to the prevailing epidemiological situations, although sentinel surveillance has now been integrated with surveillance of respiratory viruses [1]. The community-level surveillance is now focused primarily on wastewater systems as part of the public health surveillance strategy [2]. Monitoring of variants, together with environmental surveillance, is urgently warranted for future pandemic preparedness and to understand viral behavior. The Delta variant (B.1.617.2), first detected in India, evolved into a pandemic from mid-March 2021 until it was subsequently replaced by Omicron (B.1.1.529). First detected in Botswana, Hong Kong, and South Africa, the Omicron variant peaked exponentially in frequency from December 2021 to March 2022, and continued to sustain thereafter [3].

The genomic comparison of Delta and Omicron variants indicated significant mutational changes in the spike (S) protein with subsequent changes in the stability and binding to the ACE2 host receptor protein [4]. The Delta variant mainly influenced immune responses to the antigenic regions of the receptor-binding domain (RBD) of the spike protein, wherein the Omicron variant, with 30 different mutations in the spike protein, was associated with high transmissibility and vaccine-induced immune evasion [5,6]. However, the dynamics of transmission and infectivity of the Delta variant before and after the emergence of Omicron were distinct; the former was highly infectious with high rates of transmissibility, and the latter was relatively poor in the aforesaid attributes. The underlying genomic signatures and the virological and clinical significance of this difference largely remain nebulous [7–9].

Genome sequences are analyzed traditionally using alignment-based tools for the detection of mutations, level of conservation, evolutionary distances, and to assess the function of genes and proteins [10]. Viral genomes possess comparatively smaller-scale genomes, yet minimal changes result in massive functional changes in the protein [11], which are often overlooked by alignment-based tools. The lack of domain rearrangements of proteins, low sequence-alignment accuracy, especially within closely-related genomes, and incongruity for large-scale whole genome sequence, as they are memory and time-consuming [12]. In contrast, alignment-free sequence comparison methods that are based on the length and informational features of the sequences, such as base composition and quantitative distribution, have several advantages [13]. These are suitable for large-scale whole genome sequence metadata, cheaper, faster, and remain unaffected by genome recombination events. Use of machine learning methods such as k-mers in comparing SARS-CoV-2 sequences has been reported previously to analyze the variations for epidemic surveillance [14–16]. The Delta variant’s whole genome sequence data obtained as part of the national sentinel surveillance program were utilized. Here, we used k-mers and t-SNE clustering and applied a random forest classifier and a convolutional neural network for comparing sequences of Delta variants before and after the emergence of Omicron.

## 2 Materials and methods

### 2.1 Study setting and design

This observational study was conducted at the State Public Health Laboratory (Directorate of Public Health, Government of Tamil Nadu, India), which is a nodal centre for the state SARS-CoV-2 variant analysis. Since 01/03/2021, whole-genome sequencing (WGS) of SARS-CoV-2 has been conducted prospectively as part of the national and state-level genomic surveillance programs. Samples were randomly selected from the initial real-time RT-PCR–positive cases with cycle threshold (Ct) values below 25. Sequencing was performed directly from the original clinical samples using the Ion Torrent platform (Thermo Fisher Scientific, Waltham, MA, USA), following the manufacturer’s protocol. Base calling, adapter trimming, and quality control were carried out using the integrated pipeline in Ion Reporter™ Software. The processed reads were aligned to the reference genome, Wuhan-Hu-1 (GenBank accession number MN908947.3), using the IRMA assembler plugin within the Ion Reporter™ Software (Thermo Fisher Scientific, Carlsbad, CA, USA). The sequences were submitted to GenBank, and accession numbers were obtained.

Here, for this study, the sequences of Delta variants identified between 01/03/2021 and 31/12/2021, and those identified from 01/01/2022 to 31/12/2023, were retrospectively collected from the GenBank and considered as Delta variants before and after Omicron emergence, respectively. The sequences used for our study are listed in the **S1 File**. All patient information has been completely anonymized before it is accessed. The study was approved by the Institutional Ethics Committee of the Madras Medical College (EC No. 03092021).

### 2.2 Dataset description

A total of 190 genome sequences were selected, and complete S protein gene sequences were analyzed. The dataset was split into two classes, the first being the genome sequence samples collected before the omicron variant emergence, as “before Omicron”, and sequences collected after the omicron variant as “after Omicron”. All DNA sequence data were prepared in FASTA file format for further analysis. The average length of the sequences was 3813 nucleotides. From the datasets, information regarding the strain variant, the length of the sequence, and the class label was retrieved. A total of 164 sequences, as “before Omicron”, and 26 sequences as “after Omicron” were selected. As there was an imbalance of data samples between the two classes, a weighted cross-entropy loss function and random sampling were used.

This comprehensive flowchart, depicted in **Fig 1**, illustrates a sophisticated bioinformatics pipeline designed for genomic sequence analysis, encompassing three key processing stages indicated by color-coded sections. The “data input and pre-processing” section represents initial data preprocessing, where raw sequence data undergoes cleaning, standardization, and encoding into numerical representations suitable for computational analysis. The “feature extraction and exploratory data analysis” section constitutes the feature extraction and analysis phase, where k-mer frequencies are calculated alongside various statistical analyses, with pathways for both dimensionality reduction and correlation heatmaps to identify significant patterns. The machine learning model features parallel convolutional neural networks (CNN) implementations with one branch dedicated to sequence classification and another to a random forest architecture. It concludes with performance metrics, model interpretability assessments, and a comprehensive analysis [17]. This well-structured pipeline elegantly integrates sequence preprocessing, feature engineering, and advanced machine learning techniques to transform raw genomic data into actionable insights, with each component carefully designed to handle the unique challenges of high-dimensional sequence data while maximizing biological interpretation potential.

**Fig 1.**
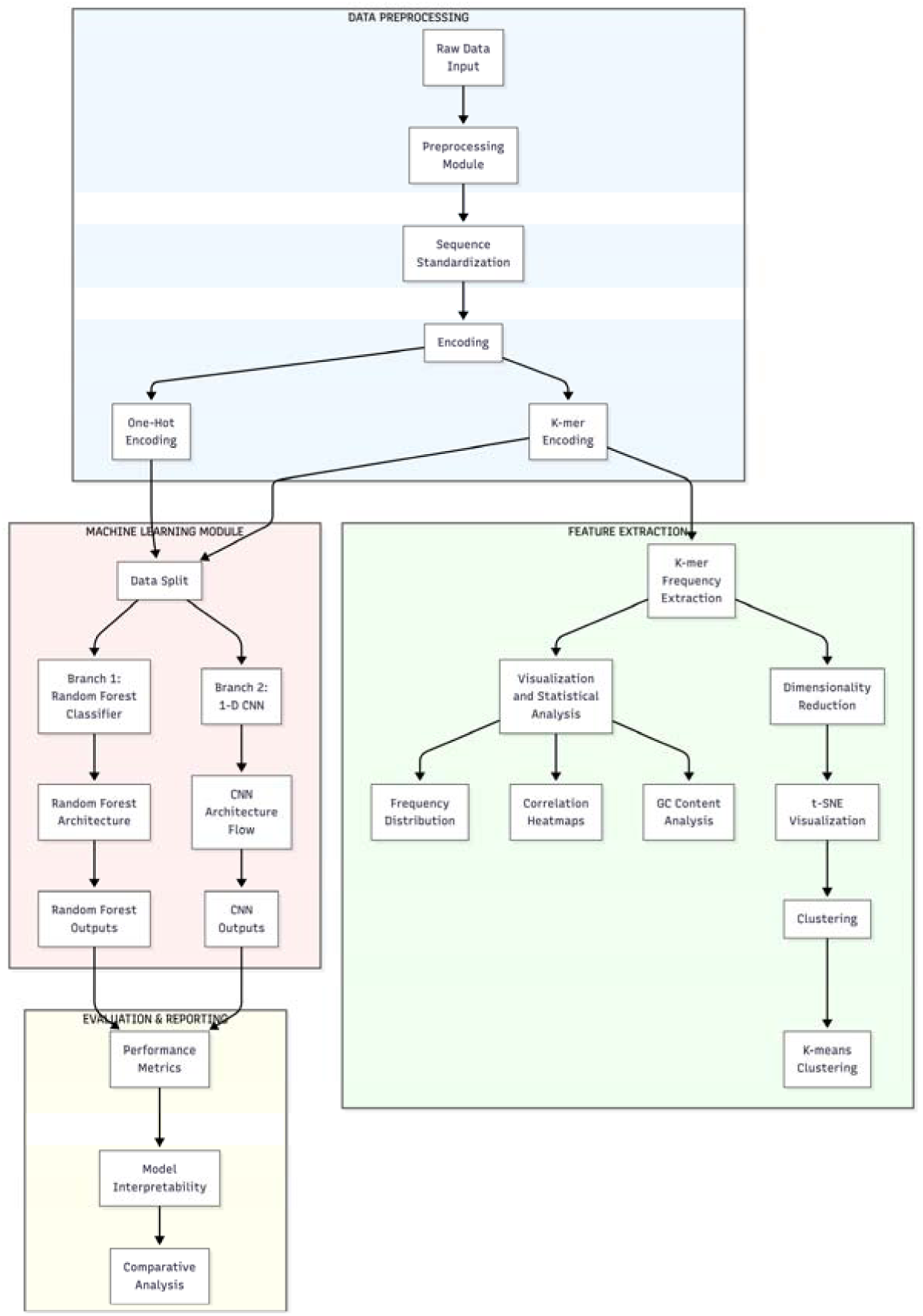
Schematic representation of the study.

### 2.3 Dataset preprocessing

The sequences were parsed using the SeqIO.parse module from Biopython (v1.84) in a Python 3 environment to read the raw FASTA files, converting the genomic data into iterable sequence records for subsequent encoding. Following sequence parsing, the data were structured into a Pandas (v2.2.2) DataFrame, where sequence data and their corresponding temporal labels (Pre-Omicron vs. Post-Omicron) were organized as discrete features, enabling efficient vectorized operations for k-mer counting and one-hot encoding. Initially, the sequences were prepared to be the same length using sequence trimming and sequence padding. With the former, the end of sequences that are larger than the smallest sequence was removed. With the latter, sequences were padded with ‘N’ characters to a fixed length of 3813 nucleotides, concerning the largest sequence length observed. Several pre-processing techniques were applied to transform DNA sequences into numerical formats suitable for DNA classification. This step was crucial, as machine learning and deep learning models required input data in numerical form.

Since DNA sequences are composed of categorical nucleotide data, they were first encoded numerically for effective classification. In this study, two encoding approaches—one-hot encoding and K-mer encoding—were utilized to achieve this transformation. Label encoding assigns a unique numerical value to each nucleotide within the DNA sequence, facilitating its use in computational models. The label encoding method assigned a numerical value to each nucleotide in the DNA sequence (A - 0, C - 0.25, G - 0.5, T - 0.75), or in one-hot encoding, the nucleotides are encoded using binary digits (A - 0001, C -0010, G - 0100, T - 1000). The DNA sequence was converted into a sequence of numbers. A “k-mer” refers to a sequence of “k” nucleotides (e.g., for k=3, the k-mers of “AGCT” would be “AGC” and “GCT”). Thus, the entire sequence was fragmented into k-mers, and every k-mer was encoded into numerical values using the encoders mentioned above. The number of unique k-mers depended on the value of k and the unique letters available in the sequence. To determine the optimal k-mer length, we conducted a sensitivity analysis across a range of k values (k=2 to 30). The results demonstrated that k=3 achieved the peak classification accuracy of 92.98% while requiring the minimum computational time (1.57 seconds). Although k=4 and k=5 yielded identical accuracy, k=3 was selected as the most efficient parameter, providing a lower-dimensional feature space that avoids the sparsity issues often associated with larger k-mer sizes. Furthermore, increasing k beyond 5 led to a noticeable decline in model performance and increased training latency, suggesting that 3-mer frequencies (representative of codon-level information) provide the most robust signal for differentiating datasets. In this study, the k value was set to 3, and the DNA sequences consisted of 4 characters. Therefore, the number of unique k-mers was equal to 4^3 or 64 k-mers. In this approach, each classifier casts a vote, and the label receiving the highest number of votes is assigned to the sample, as illustrated in **Figs 2A and 2B**.

**Fig 2.**
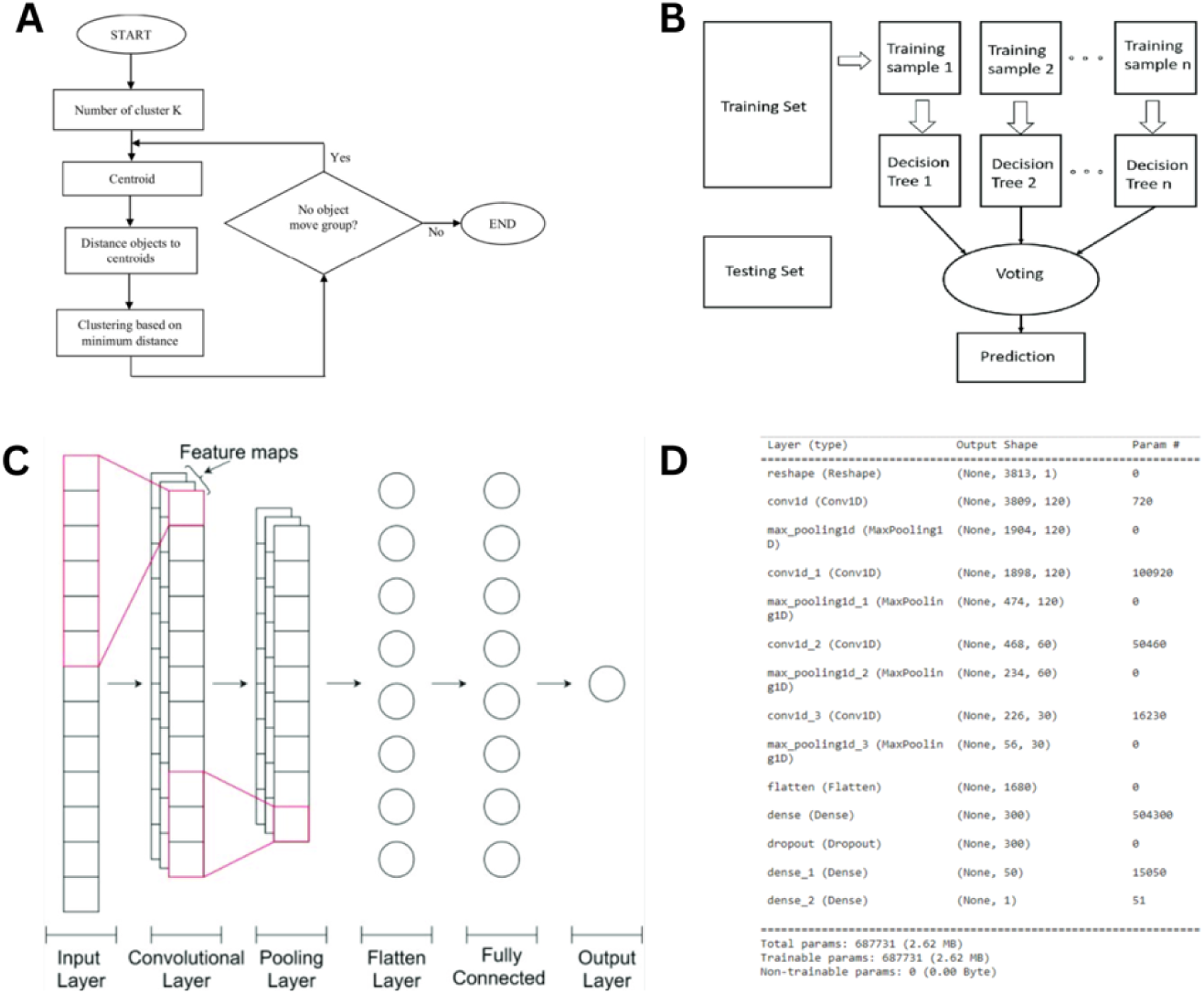
Machine Learning Model Architectures and Workflow for Genome Classification. (A) k-means clustering algorithm workflow (B) Schematic architecture of the random forest classification model (C) Architecture of the 1-D convolutional neural network for whole-genome sequence classification. (D) Layer-wise structural summary of the 1-D CNN model configuration

### 2.4 Exploratory data analysis

The preprocessed data was further analyzed for underlying patterns using various tools. The k-mer frequency in each DNA sequence was calculated using the Count Vectorizer function from the Scikit-learn library. The frequency of k-mers is used as a feature in machine learning models. The correlation between each k-mer was plotted in the graph to analyze the relation between the k-mers.

### 2.5 T-distributed stochastic neighbor embedding

t-SNE was applied for dimensionality reduction of high-dimensional genomic feature vectors to visualize sample clustering in two- and three-dimensional space. The dimensionality reduction to two components compressed the data’s inherent structure, bringing the clusters closer together while maintaining their distinctiveness. Both clusters exhibit remarkably similar internal cohesion, with Class 0 showing a dispersion of 11.41 and Class 1 showing 11.95, indicating consistent within-class variability. The variance is almost equally distributed between the two components, with Component 1 accounting for 51.79% and Component 2 for 48.21% of the total point distribution. This balanced contribution from both dimensions indicated that the 2D representation effectively captured the underlying structure of the data, with neither component dominating the visualization, making this a reliable low-dimensional representation for further analytical interpretation.

### 2.6 Machine learning models

Random forest and 1D CNN models were used to analyze genomic sequences. The random forest classifier generated predictions using multiple decision trees and majority voting. The 1D CNN model processed full-length sequences through convolutional layers to learn discriminative patterns. Both models were trained and evaluated on designated datasets.

### 2.7 K-means clustering

K-means Clustering, as an unsupervised machine-learning method, was implemented to identify the shift of clusters over a period of time to identify the genomic sequences for specific mutation patterns. K-means clustering was applied to group sequences into K clusters based on similarity. The algorithm iteratively assigned samples to the nearest centroid and updated cluster centers until stable groupings were obtained [18,19].

### 2.8 Random forest classifier

For classification, a random forest model was implemented as an ensemble learning approach. The algorithm constructs multiple decision trees using bootstrap-sampled subsets of the training data and random subsets of features at each split. Each tree independently predicts a class label for a given input sample. Final classification was determined using majority voting across all trees. This procedure was applied to assign labels to previously unlabeled samples. In this study, the random forest model is trained with 80% of the dataset, and 20% of the dataset is used to test the model. It operates by selecting random subsets of data and features for each tree, ensuring diversity among the trees. The Random Forest model functions as an ensemble of N independent decision trees. By iteratively testing values for N (number of estimators) ranging from 10 to 5,000, we determined that N=1000 provided optimal performance, ensuring a sufficient diversity of trees for robust majority voting without incurring unnecessary computational or memory overhead. Each branch within these trees represents a binary split based on the normalized frequency of a specific 3-mer pattern. By traversing these branches, the model partitions the genomic data into increasingly homogeneous subsets, eventually reaching a leaf node that assigns the sequence to either the before or after Omicron emergence group. Each decision tree in the forest splits nodes based on a chosen feature and threshold that best separates the data, typically using criteria like Gini impurity or entropy in classification tasks and mean squared error (MSE) in regression tasks. Gini Impurity and Entropy are two key criteria used in decision trees to measure the quality of splits. Both metrics evaluate how “pure” a node is, meaning how mixed the classes are in a classification problem. A lower value for either metric indicates a more homogeneous (pure) node.

### 2.9 Gini impurity calculation

The probability of the sample being incorrectly classified is labeled based on the distribution of classes within a node was calculated by Gini impurity using the formula:

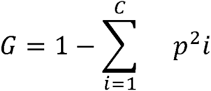

Where C is the number of classes, and p is the proportion of samples that belong to class i at the node.

### 2.10 Entropy Calculation

To quantify the amount of uncertainty in a node, entropy was calculated using the formula:

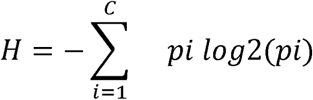

where H is the entropy, C is the number of classes, and p is the probability of class i in the node. Entropy is 0 for a pure node and reaches its maximum when all classes are equally probable, meaning the node is most uncertain.

### 2.11 1-D convolutional neural network

In addition to using a random forest to identify critical k-mers that distinguish genome sequence classes, we implemented a 1-D CNN to operate directly on individual nucleotides rather than k-mers. **Fig 2C** represents the architecture of a 1-D CNN model. Neural networks are made of layers and nodes, where each layer is designed to perform a specific task(s). The CNN consisted of convolutional layers where each layer performed the convolution operation on its input data. A convolution layer computes its output by convolving the input with its filter weights, adding a bias β, and passing each result through an activation function. In 1D convolution, for *N* number of kernels or filters, a stride of one, and a dilation rate of one, the i^th^ hidden unit h_i_ can be expressed as:

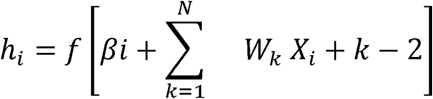

Here, *f* [.] corresponds to the activation function, βi is the bias, and *w* and *x* are the weights and data points, respectively. In other words, a 1D CNN slides a filter along one dimension, computing the dot product of the weight and input data, and creates the output feature map of the convolution layer. The model included the input layer, convolutional layers, a flatten layer, a dense layer, and an output layer. The network had a total of 687,731 trainable parameters. The model summary is shown in **Fig 2D**.

In this study, TensorFlow (v2.x) was utilized to construct the Deep Learning pipeline. The model architecture begins with a Reshape layer to organize the input genome sequences into a three-dimensional format suitable for convolution. Subsequently, three 1D convolutional layers, each followed by max-pooling, extract hierarchical features from the sequence while reducing dimensionality. A Flatten layer then converts the resulting feature maps into a one-dimensional vector. This vector is fed into two Dense layers for high-level feature combination, interspersed with a Dropout layer to mitigate overfitting. Finally, a single-node output layer provides the classification outcome. The combination of convolution, pooling, and dense layers yields an end-to-end architecture optimized for genomic data. The model was compiled using the Adam optimizer, which adaptively updates network weights by computing individual learning rates for each parameter based on the first and second moments of the gradients. This approach helps the network converge more quickly and reliably compared to standard stochastic gradient descent. A binary cross-entropy loss function was employed to measure the discrepancy between predicted probabilities and actual labels, making it well-suited for binary classification tasks.

The training dynamics of the 1D-CNN revealed a non-monotonic learning trajectory characteristic of deep representation learning in high-dimensional genomic data. The model was trained for a maximum of 100 epochs, utilizing an initial learning rate of 1e-3 with a scheduler configured to decay the rate to a minimum floor of 1e-6. The model underwent an initial phase of high variance (Epochs 1–15), where validation loss exhibited significant fluctuations (peaking at 0.71). This volatility indicates the model was actively exploring the optimization landscape to escape sharp local minima rather than prematurely converging on suboptimal solutions. Following this exploration, the model entered a stabilization phase where the validation loss decreased and plateaued, signaling a shift from memorizing noise to learning robust, generalizable genomic motifs. ReduceLROnPlateau monitored validation performance and systematically reduced the learning rate when progress plateaued, allowing the model to fine-tune its parameters more effectively and avoid suboptimal convergence. Although local minima were observed in early epochs lacked stability, Epoch 33 was identified via Early Stopping as the optimal convergence point. At this epoch, the model achieved a critical equilibrium, minimizing training loss (0.598) while maintaining a stable validation loss, thereby ensuring that the classification relied on learned structural features rather than stochastic fluctuations.

### 2.12 Metrics

#### 2.12.1 Accuracy

Accuracy measures the proportion of correctly classified instances among all predictions, providing an overall gauge of a model’s performance. Accuracy is calculated using the following formula:

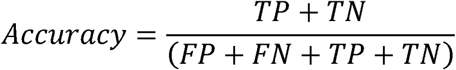

True positives (TP) refer to instances in which the model correctly identifies the positive class, while true negatives (TN) indicate cases where the model accurately predicts the negative class. False positives (FP) occur when the model incorrectly classifies a negative instance as positive, and false negatives (FN) arise when a positive instance is mistakenly categorized as negative. Together, these values provide a comprehensive picture of classification performance.

#### 2.12.2 F1 score

F1 score is the harmonic mean of precision and recall, offering a balance between these two complementary metrics. The F1 score provided an informative single value that captured both how precise the model’s positive predictions are and how well the model detected all actual positive [20,21]. As such, it was particularly useful in applications where one aims to balance false positives and false negatives, especially in the presence of class imbalance. The F1 score is defined as:

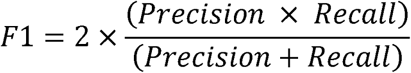

Precision and recall are two key metrics used to evaluate a classifier’s performance on a positive class. Precision measures the proportion of correctly identified positives among all instances labeled as positive, reflecting how reliable the model’s positive predictions are. Formally, it is given by

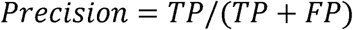

Recall, on the other hand, captures the proportion of actual positives that the model successfully detects. It is calculated as

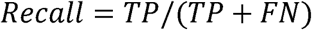

Together, precision and recall provide complementary insights into a model’s accuracy in identifying positive instances.

#### 2.12.3 SHapley Additive exPlanations

SHAP (SHapley Additive exPlanations) is a method grounded in cooperative game theory that provides individualized explanations for model predictions. It assigns each feature a “Shapley value” indicating the contribution of that feature to the prediction relative to a baseline [22]. For a model *f* and a set of features *F*, the SHAP value *φ* for feature *i* is computed by averaging the difference in model predictions with and without the feature, over all possible feature subsets, *S* ⊆ (*F* ∖ {*i*}). Formally:

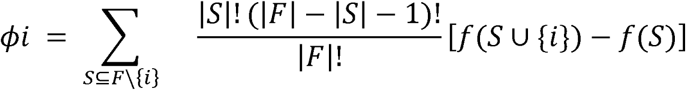

Where ∣*F*∣ is the total number of features, and *f*(*S*) denotes the model’s prediction when only the features in *S* are present.

SHAP values are based on a rigorous game-theoretic foundation, and uphold desirable properties such as local accuracy, consistency, and missingness. SHAP was used to generate both global and local explanations: globally, by aggregating contributions across many samples to understand how features influence overall model behavior, and locally, by explaining a single instance’s prediction in terms of its feature contributions.

### 2.13 Receiver operating characteristic

The ROC curve was used to evaluate the performance of binary classification models by plotting the True Positive Rate (TPR) against the False Positive Rate (FPR) at various decision thresholds. The TPR represents the proportion of actual positives correctly identified, while the FPR measures how many negative instances are incorrectly classified as positive. By adjusting the classification threshold, from classifying almost everything as positive to classifying almost everything as negative, the ROC curve was plotted to reveal the trade-off between detecting more true positives and avoiding false positives. A model whose curve hugs the top-left corner of the plot (high TPR, low FPR) is considered superior, as it consistently distinguishes between classes across a range of thresholds. The Area Under the Curve (AUC) condenses this information into a single metric, with higher values (closer to 1.0) signifying better overall discrimination.

## 3 Results

SARS-CoV-2 Delta variant spike gene sequences identified during 2021 and 2022 outbreaks were investigated for genomic differences between the strains identified before Omicron and after Omicron emergence. In this study, the sequences were analyzed using statistical methods and machine-learning models. The models were trained and fitted with the dataset such that the accuracy is maximized. The primary goal of the study was to understand and find the difference in DNA sequences. The machine learning models were built to classify the classes of sequences and then retrieve the features that provide a significant contribution to the classification. The temporal trend suggested an overall number of point mutations (n=157) from both groups of data, suggesting stable changes in genomic sequence that had occurred over time. The comparison of point mutations displayed four significant mutations that disappeared in the “after Omicron” sequences compared to the “before Omicron” sequences.

**Fig 3A** shows the list of k-mers used in this study. The frequency of k-mers was used as a feature in machine learning models. The frequency distribution of every k-mer across the dataset is shown in **Fig 3B**. The correlation heat of the k-mers is shown in **Fig 3C**. Blue denotes a negative connection, whereas red denotes a positive correlation. GC content denotes the proportion of guanine (G) and cytosine (C) nucleotides within a DNA or RNA sequence and serves as a critical parameter in genomic analysis. The GC content of the sequences was analyzed, and the GC percentage was plotted for “before Omicron” (**Fig 3D)** and “after Omicron” (**Fig 3E**). Both plots show that the mean GC content remains consistent between before Omicron and after Omicron sequences, suggesting no significant shift in overall GC composition. This implies that any observed differences between these two classes are likely influenced by other genomic features or sequence variations, rather than changes in GC content alone.

**Fig 3.**
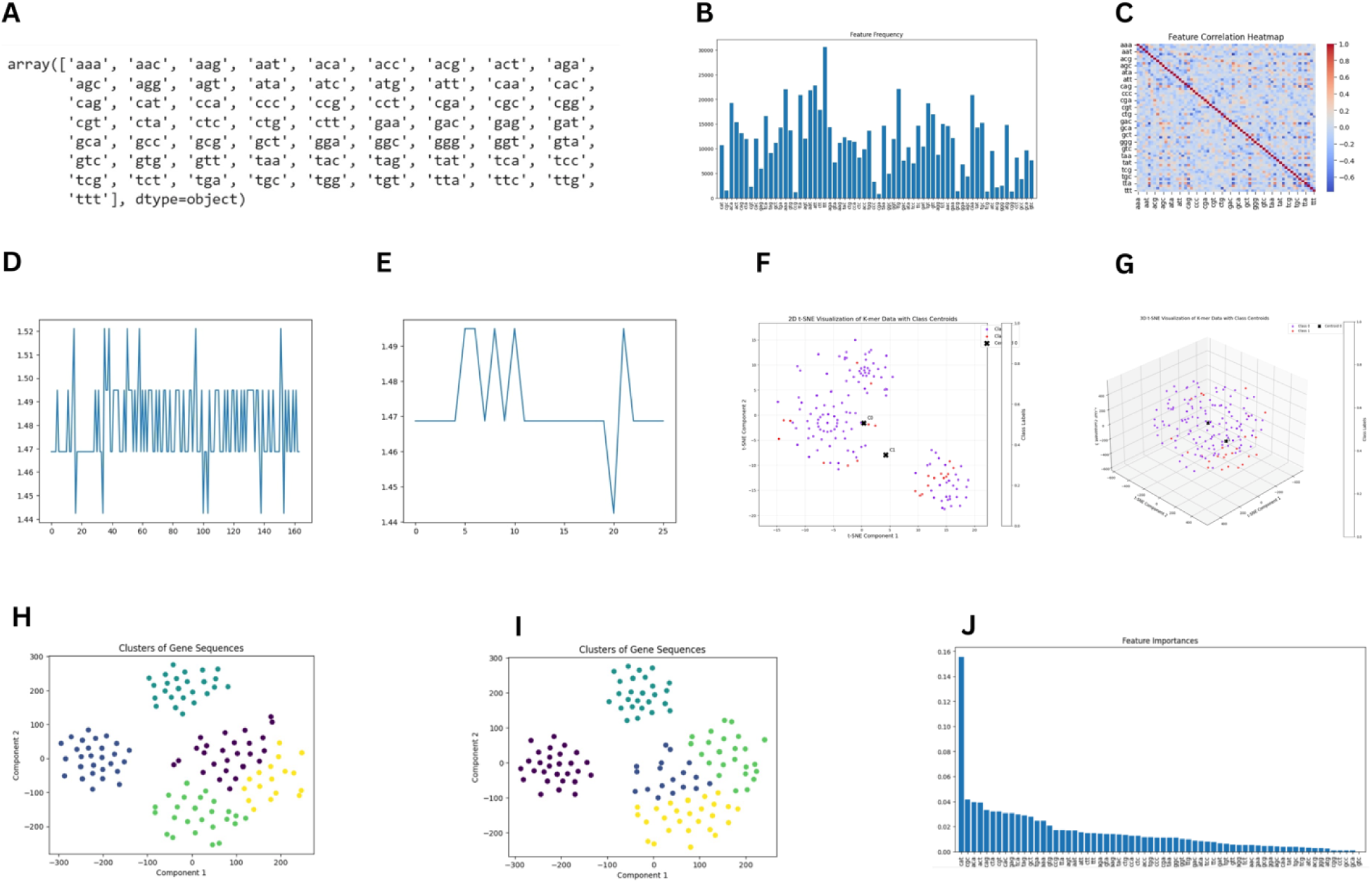
K-mer–Based Feature Analysis and Clustering of Delta Variant Genomes Pre- and Post-Omicron Emergence. (A) Comprehensive set of all possible 3-mer sequence combinations used for feature extraction. (B) Global frequency distribution of k-mers across the entire sequence dataset. (C) Correlation heatmap depicting interrelationships among k-mer frequencies. (D) GC content distribution in Delta variant genomes prior to Omicron emergence. (E) GC content distribution in Delta variant genomes following Omicron emergence. (F) Two-dimensional t-SNE projection illustrating clustering patterns of DNA sequences. (G) Three-dimensional t-SNE visualization demonstrating class separation in sequence data. (H) K-means clustering of Delta variant sequences before Omicron emergence. (I) Cluster structure revealed by K-means partitioning of pre-Omicron Delta genomes. (J) Random forest–derived feature importance scores highlighting key predictive k-mers

The 2D t-SNE visualization shown in **Fig 3F** revealed a moderate separation between the two classes, with a centroid distance of 7.49 units—significantly smaller than observed in the 3D representation. Based on the t-SNE visualization results in 3D space, as shown in **Fig 3G**, the data revealed distinct clustering patterns with meaningful separations between classes. The distance matrix shows a substantial separation of 256.72 units between cluster centroids, indicating well-differentiated groupings despite some internal variation. Both clusters display relatively high dispersion values (350.43 for Class 0 and 369.13 for Class 1). The distribution of variance across the three components was balanced, with Component 3 capturing the highest proportion at 40.45%, followed by Component 2 (34.94%) and Component 1 (24.61%). The balanced distribution indicated that three-dimensional representation effectively captures the complex relationships in the data, with information meaningfully distributed across all dimensions rather than being dominated by a single component.

This compression effect demonstrated how the additional dimension in 3D space allowed for a more expansive distribution of data points, potentially revealing subtleties in genetic relationships that become condensed in 2D.

The cluster analysis collected from the K-Means classification results indicates stable evolution with some significant changes over time (**Fig 3H and 3I**), which showed some mutations moving within the class and some mutations classifying into different classes in the “after Omicron” Delta variant classification result. Clusters in before-omicron Delta variants are shown in **Fig 3H**, and Clusters in after-omicron Delta variants in **Fig 3I**.

In our study, we found 157 ‘persistent’ mutations identified from both before and after the omicron group of delta variants and four ‘vanished’ mutations (C2011G, C643T, T776A, C179T) that were lost after the omicron group of delta variants. Among the separated clusters, cluster shifts were identified between and within the clusters **(Fig 3H and Fig 3I)** due to genomic feature differences and includes (4,4):29; (2,2):33; (4,0):14; (3,3):32; (1,0):3; (1,4):12; (1,3):3; and (0,1):38.

The random forest classifier was trained on 80% of the dataset, using 1000 trees (estimators) as the main function parameter, resulting in a 93% accuracy and F1 scores of 96% and 75% for different classes. This high accuracy and balanced F1 metric indicate robust performance across classes. Furthermore, the default random forest feature importance calculation (often based on Gini importance) was used to quantify the contribution of each k-mer to the classification, and these importances were subsequently plotted to visualize the most influential features (**Fig 3J**).

The importance of the feature was calculated using the SHAP values (Shapely additive explanation). A positive SHAP value indicates that the feature increases the prediction (e.g., pushes the model towards a positive class in classification). A negative SHAP value indicates that the feature decreases the prediction (e.g., pushes the model away from a positive class). As shown in **Fig 4A**, k-mers “CAT”, “ACA”, and “CTA” had the highest SHAP value. Whereas k-mer “CAT”, “CGC”, and “ACA” had the highest importance according to the random forest classifier.

**Fig 4.**
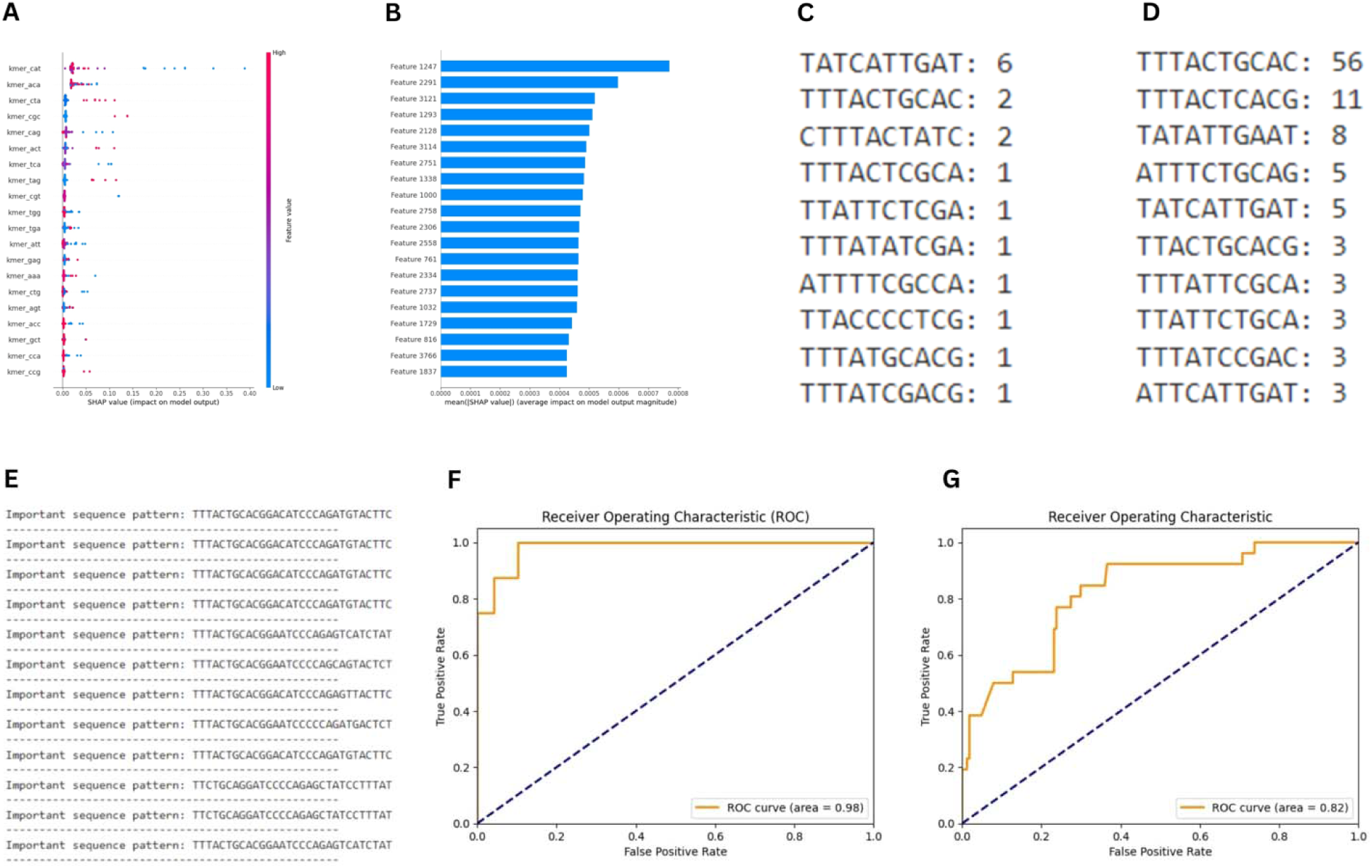
Model Interpretation, Motif Discovery, and Classification Performance Analysis. (A) Comparative k-mer importance profiles derived from SHAP attribution and random forest metrics. (B) Genome-wide positional SHAP attribution landscape showing nucleotide-level contributions to model predictions. (C) Most prevalent sequence motif in pre-Omicron Delta genomes with frequency counts. (D) Most prevalent sequence motif in post-Omicron Delta genomes with frequency counts. (E) Dataset-wide discriminative motif patterns learned by the 1-D CNN model. (F) Receiver operating characteristic curve of the random forest model illustrating threshold-dependent classification performance. (G) Receiver operating characteristic curve of the 1-D CNN model depicting sensitivity–specificity trade-off

The custom 1-D CNN model took an input of shape (3813, 1) and had multiple 1-D CNN layers with dense layers to classify the sequences. The 1-D CNN model produced an accuracy of 77% and an F1 score of 86% and 36%, as shown in **Table 1**. This approach provided a more granular view of the sequence, as convolutional filters captured local motifs and patterns in an end-to-end manner without explicit feature engineering. By comparing the random forest’s k–mer–based feature importance with the 1-D CNN’s learned representations, evaluation of differences in predictive performance and interpretability was possible. The SHAP values were calculated for each position of the sequence since the 1D-CNN model took the entire genome sequence without converting it into k-mers as input. The nucleotide position is shown in **Fig 4B**.

**Table 1.** Comparison of the random forest model and the 1-D CNN model in classifying the genome sequences.

| Model | Class | Precision | Recall | F1-score | Support | Accuracy |
| --- | --- | --- | --- | --- | --- | --- |
| Random Forest Classifier | 0 | 0.96 | 0.96 | 0.96 | 49 | 0.93 |
|  | 1 | 0.75 | 0.75 | 0.75 | 8 |  |
| 1-D CNN | 0 | 0.93 | 0.80 | 0.86 | 189 | 0.77 |
|  | 1 | 0.27 | 0.54 | 0.36 | 26 |  |

The most commonly occurring pattern in the Delta variant “before Omicron,” along with its count, is shown in **Fig 4C**, and the most commonly occurring pattern “after Omicron along with its count, is shown in **Fig 4D**. The important patterns in the DNA sequences were recorded based on the 1-D CNN model. These patterns provided the difference between the two classes. **Fig 4E** shows the important patterns in the entire dataset. Receiver Operating Characteristic (ROC) curves of the random forest classifier model and 1-D CNN model, respectively, are shown in **Figs 4F and 4G**. The ROC curve was used for graphical representation to evaluate the performance of a binary classification model. It plotted the True Positive Rate (TPR) against the False Positive Rate (FPR) at various threshold settings. The curve showed the trade-off between sensitivity (TPR) and specificity (1 - FPR) across different thresholds. A model with a curve closer to the top-left corner indicated better performance.

## 4 Discussion

A diverse group of RNA viruses and their ability to constantly mutate during their evolutionary process are too complicated to comprehend and prepare for future pandemics [23]. The availability of next-generation sequencing data has provided new opportunities and achievements in the study of viral evolution and ecology, as well as challenges in handling the huge volume of raw sequence data [24]. Viral sequences are regularly monitored for genomic and proteomic data analysis, such as mutational and structural changes, using available bioinformatic tools. Such analyses are useful for studying its phenotypic features, drug discovery, epidemiological investigations, identifying new variants, and their clinical implications [22,25,26]. However, such alignment-based methods do not provide sufficient information due to their inherent limitations [27].

Previously, studies on sequence classification on the SARS-CoV-2 spike gene have been carried out using a k-mer-based approach for training the data to identify a specific variant [28]. The k-mer extraction, as it does not depend on sequence homology, is considered more effective in classifying distant or closely related strains as well as for large sequencing datasets, and possesses high computational speed, memory efficiency, and biological functionality. In our study, the model trained had high accuracy, as an effective feature of the genome sequence was extracted. The model was trained to gain a temporal trend suitable for the viral sequences of a specific variant that emerged at different periods.

Alignment-free sequence analyses are used to study the evolution of microorganisms and cis-regulatory molecules, and to analyze meta information such as next-generation sequencing data [29,30]. Viral sequences have been analyzed using an ML approach through mapping of the sequence in a feature space, followed by data processing using ML techniques. These methods have been shown to be highly valuable in classifying different viruses, including dengue viruses, HIV-1, influenza A, hepatitis B, and hepatitis C [31–33].

GC content variations can reveal structural and functional regions within genomes, such as promoter sites or regions prone to mutation, aiding in both annotation and comparative genomics studies [34]. The SHAP values provide a way to explain the contribution of each feature to a specific prediction, making models more transparent and understandable, especially in complex models like deep learning [35,36]. The 1D CNN approach provided a more granular view of the sequence, as convolutional filters captured local motifs and patterns in an end-to-end manner without explicit feature engineering. By comparing the random forest’s k–mer–based feature importance with the 1-D CNN’s learned representations, evaluation of differences in predictive performance and interpretability was possible. Elevated GC content is typically associated with enhanced stability of the nucleic acid duplex, owing to the stronger triple hydrogen bonding between G and C bases, in contrast to the double hydrogen bonds formed between adenine (A) and thymine (T). This property can influence the melting temperature of DNA and the efficiency of PCR, making it a crucial factor in experimental design [37].

All sequences included in this study were confirmed to belong exclusively to the B.1.617.2 lineage based on their original annotations and verification prior to analysis, with no A.Y.* sublineages present. Thus, the “before Omicron” and “after Omicron” groups represent samples from different time periods within the same lineage, rather than different Delta sublineages. The classification differences detected by the k-mer–based models therefore reflect subtle genomic changes over time within B.1.617.2, rather than shifts in lineage assignment.

Following the identification of specific mutations within the “before Omicron” class and specific mutations that disappeared in the “after Omicron” class, the random forest classifier model was trained with 1000 samples, providing an accuracy of 93%, and F1 scores of 96% and 75%. This ensured that the model classified the dataset with high accuracy, and the importance factor of each feature (k-mer) was extracted using the default method of the random forest classifier [13]. Following the identification of specific mutations present in the “before Omicron” class and those that disappeared in the “after Omicron” class, we trained a random forest classifier on 1000 representative samples. The model achieved a high overall accuracy of 93%, reflecting its strong predictive capability in distinguishing between the two temporal variant classes. The F1 score for the majority class was 96%, indicating excellent precision and recall, while the minority class yielded an F1 score of 75%, which is still acceptable given the class imbalance. These results underscore the robustness of the classifier in capturing subtle mutational patterns associated with variant transitions. Additionally, we extracted the importance scores of each k-mer feature using the built-in feature ranking method of the random forest algorithm, enabling interpretability and highlighting the most influential mutations contributing to classification performance. This provides valuable insight into evolutionary shifts in the viral genome, particularly during the emergence of Omicron and its impact on the mutational landscape.

In this study, the sequences were analyzed using statistical methods and machine-learning models. The models were trained and fitted with the dataset such that the accuracy is maximized. The machine learning models were built to classify the classes of sequences and then retrieve the features that provide significant contributions to the classification. The Temporal Trend report suggested an overall number of 157 persistent point mutations from both groups of data, suggesting stable changes in genomic sequence over time. The comparison of point mutations displayed four significant mutations that vanished in the “after Omicron” sequence data. The cluster analysis derived from the K-Means classification results indicates stable evolution with some significant changes over time, which showed some mutations moving within the class and some mutations classifying into different classes in the “after Omicron” Delta variant classification. The k-mer-based supervised classification approach presented in this paper offers several advantages over commonly used software tools for virus subtype classification. Our evaluations on multiple manually curated datasets demonstrate that k-mer classification enables rapid and accurate SARS-CoV-2 subtyping, outperforming many current state-of-the-art methods.

This study has several limitations that should be considered when interpreting the findings. The sample size was modest (n = 190), with a marked class imbalance between pre-Omicron and post-Omicron groups, which may influence classifier generalizability despite the use of weighting strategies. In this study, alignment-free approaches were selected to explore complementary analytical capabilities, particularly for pattern recognition and machine-learning–driven classification, and scalability advantages become relevant primarily in larger datasets.

Only spike gene sequences were analyzed rather than complete genomes; therefore, conclusions reflect spike-specific evolutionary patterns and may not represent genome-wide dynamics. The study design was retrospective and observational; thus, associations identified between temporal groups and sequence features should not be interpreted as causal evolutionary drivers. Model evaluation was performed on a single dataset without external validation, which limits inference regarding performance on independent or future datasets. Because k-mer frequencies summarize sequence content, convergent compositional similarity could theoretically yield classification signals unrelated to true evolutionary relatedness. Taken together, these considerations indicate that while the alignment-free machine learning framework used here is effective for detecting discriminative sequence patterns, it does not replace traditional alignment-based genomic analyses. Instead, each approach provides distinct analytical advantages, and integrative strategies combining positional mutation analysis with compositional feature learning may offer the most comprehensive framework for future viral evolutionary studies [38].

## 5 Conclusions

This study offers valuable insights into the temporal evolution of SARS-CoV-2 Delta variants in the context of the emergence of Omicron. Identification of persistent and vanished mutations among the delta variants, possibly influenced by the emergence of Omicron variants, contributes to the understanding of viral genomic stability and adaptability. The integration of alignment-free methods and machine learning models (e.g., K-means and random forest) provides an effective approach for uncovering subtle yet meaningful genomic changes that may not be apparent through traditional sequence comparisons. The ability to classify variants and identify influential k-mers enhances molecular surveillance capabilities, potentially aiding in early detection of functionally significant mutations. These findings may support public health efforts in tracking the evolution of variants, inform adjustments to vaccine strategies, and guide future studies in viral pathogenesis and evolutionary modelling.

## Acknowledgements

ML is supported by grants through AI52731, the Swedish Research Council, the Swedish Physicians against AIDS Research Foundation, the Swedish International Development Cooperation Agency, SIDASARC, VINNMER for Vinnova, Linköping University Hospital Research Fund, CALF, and the Swedish Society of Medicine. VV is supported by: The NIH Office of Research Infrastructure Programs (P51 OD011132 to ENPRC), and Emory CFAR (P30 AI050409). The authors thank all the national and international members of the Infectious Diseases Society of India (IDSI), [https://idsi.org.in/], Chennai, for extending insightful discussions as well as technical and logistic support.

## Authors contribution

Authors SS, DJ, PB, ML, VV, RS, and EMS were involved in conceptualization, study design, and supervision. Authors KA and PS were involved in data collection, analysis, interpretation, and manuscript preparation. Author AJ was involved in supervision, data analysis, and interpretation. All authors revised and approved the submitted version of the manuscript.

## Declarations

### Data availability statement

All the data generated in the study are presented in the manuscript, illustrations, and supplementary data.

### Ethics approval and consent to participate

The protocols involving human participants were approved by the Institutional Ethics Committee of the Madras Medical College (EC No. 03092021).

### Clinical trial registration

Not applicable

## Supplementary Files

**S1 File. GenBank accession numbers of the sequences analyzed**

